# Loss of Activity-Induced Mitochondrial ATP Production Underlies the Synaptic Defects in a *Drosophila* model of ALS

**DOI:** 10.1101/2021.12.14.472444

**Authors:** Nicholas E. Karagas, Kai Li Tan, Hugo J. Bellen, Kartik Venkatachalam, Ching-On Wong

**Affiliations:** Department of Integrative Biology and Pharmacology, McGovern Medical School at the University of Texas Health Sciences Center (UTHealth), Houston, TX 77030, USA; Departments of Molecular and Human Genetics and Neuroscience; Graduate Program in Developmental Biology, Baylor College of Medicine, Houston, TX 77030, USA; Duncan Neurological Research Institute, Texas Children Hospital, Houston, TX 77030, USA; Graduate Program in Biochemistry and Cell Biology MD Anderson Cancer Center and UTHealth Graduate School of Biomedical Sciences; Graduate Program in Neuroscience MD Anderson Cancer Center and UTHealth Graduate School of Biomedical Sciences; Department of Biological Sciences, Rutgers University, Newark, NJ 07102, USA; Department of Neurology, University of Washington, Seattle, WA 98195, USA; Emergent BioSolutions, Gaithersburg, MD 20879, USA

**Keywords:** *Drosophila* neurobiology, neurodegeneration, ALS, VAPB, ER Ca^2+^ signaling, mitochondrial ATP production, activity-induced ATP production, neuronal bioenergetics

## Abstract

Mutations in the gene encoding VAPB (vesicle-associated membrane protein B) cause a familial form of Amyotrophic Lateral Sclerosis (ALS). Expression of an ALS-related variant of *vapb* (*vapb^P58S^*) in *Drosophila* motor neurons results in morphological changes at the larval neuromuscular junction (NMJ) characterized by the appearance of fewer, but larger, presynaptic boutons. Although diminished microtubule stability is known to underlie these morphological changes, a mechanism for the loss of presynaptic microtubules has been lacking. Here, we demonstrate the suppression of *vapb^P58S^*- induced changes in NMJ morphology by either the loss of ER Ca^2+^ release channels or the inhibition Ca^2+^/calmodulin (CaM)-activated kinase II (CaMKII). These data suggest a model in which decreased stability of presynaptic microtubules at *vapb^P58S^* NMJs result from hyperactivation of CaMKII due to elevated cytosolic [Ca^2+^]. We attribute the Ca^2+^ dyshomeostasis to delayed extrusion of cytosolic Ca^2+^ stemming from a paucity of activity-induced mitochondrial ATP production coupled with elevated rates of ATP consumption. Taken together, our data point to bioenergetic dysfunction as the root cause for the synaptic defects in *vapb^P58S^*-expressing *Drosophila* motor neurons.

**Significance Statement:** Rates of ATP production and consumption are tightly synchronized in healthy neurons. Whether this synchrony is lost in models of neurodegenerative diseases remains poorly understood. Here, we find that expression of a gene encoding an ALS-causing variant of an ER membrane protein, VAPB, decouples mitochondrial ATP production from neuronal activity. Due to a combination of diminished ATP production and elevated ATP consumption — established outcomes in ALS neurons — the levels of ATP in *vapb^P58S^* neurons are unable to keep up with the bioenergetic burden of depolarization. The resulting paucity of ATP and attendant decrease in the activity of Ca^2+^ ATPases results in diminished extrusion of cytosolic Ca^2+^ in *vapb^P58S^*-expressing motor neurons. The accumulation of residual Ca^2+^ in *vapb^P58S^*-expressing neurons underlies paired-pulse facilitation of synaptic vesicle release, and the changes in bouton development at the NMJ. In summary, our findings point to bioenergetic dysfunction due to the loss of activity-induced ATP production as being the cause of the synaptic defects observed in a *Drosophila* model of ALS.

## Introduction

ALS is an untreatable neurodegenerative disease characterized by the progressive loss of motor function leading to paralysis and death by respiratory failure (Taylor et al., 2016). Mutations in many genes have been implicated in the onset of the familial forms of the disease, which result in one or more of the following — disrupted nucleocytoplasmic transport, proteostatic imbalance, alterations in RNA metabolism, genome instability, mitochondrial dysfunction, aberrant Ca^2+^ homeostasis, neuronal hyperexcitability, and neuroinflammation (Ling et al., 2013; Selfridge et al., 2013; Taylor et al., 2016; Lin et al., 2017; Frere and Slutsky, 2018). In this study, we sought to examine the mechanisms of neuronal dysfunction in flies expressing a missense variant of *vapb*, which is equivalent to the human variant that causes a familial form of ALS (ALS8) in humans (Nishimura et al., 2004, 2005; Kanekura et al., 2006; Marques et al., 2006; Landers et al., 2008; Borgese et al., 2021). We chose VAPB because of the importance of this protein to neuronal viability. In addition to the missense variant that functions via a dominant-negative mechanism (Ratnaparkhi et al., 2008), spinal cords from patients with sporadic cases of ALS exhibit decreased *VAPB* expression (Anagnostou et al., 2010; Mitne-Neto et al., 2011). Defective VAPB function is also observed in Parkinson’s disease (Kun-Rodrigues et al., 2015; Paillusson et al., 2017; Boczonadi et al., 2018).

At the molecular level, VAPB is a single-pass ER membrane protein that has been proposed to regulate mTOR signaling, autophagy, lysosomal acidification, proteasomal degradation and ER quality control, and formation of interorganellar contacts that link the ER to endosomes, peroxisomes, Golgi, and mitochondria (Peretti et al., 2008; De Vos et al., 2012; Deivasigamani et al., 2014; Moustaqim-Barrette et al., 2014; Roulin et al., 2014; Dong et al., 2016; Stoica et al., 2016; Hua et al., 2017; Zhao et al., 2018; Chaplot et al., 2019; Mao et al., 2019; Ş türk et al., 2019; Borgese et al., 2021). By en promoting the formation of ER–mitochondria contacts, VAPB also mediates the transfer of Ca^2+^ from ER into the mitochondrial matrix, and thereby, regulates ATP synthesis (De Vos et al., 2012; Stoica et al., 2016; Gomez-Suaga et al., 2017; Smith et al., 2017; Xu et al., 2020; Borgese et al., 2021; Wong et al., 2021).

Expression of the ALS-related variants of *vapb* in *Drosophila* neurons have been shown to result in morphological alterations in both axons and dendrites (Chai et al., 2008; Ratnaparkhi et al., 2008; Kamemura et al., 2021). Although it is known that defects in the development of the axon termini stem from diminished stability of presynaptic microtubules (Chai et al., 2008; Ratnaparkhi et al., 2008), exactly how mutant VAPB disrupts the microtubule cytoskeleton has remained unknown. In order to uncover the underlying molecular mechanism, we sought to examine genetic interactions between the *vapb* variant and genes encoding ER Ca^2+^ channels whose absence elicits phenotypes that ostensibly resemble those evoked by mutant *vapb* (Wong et al., 2014). These studies revealed that the phenotypes arising from the expression of mutant *vapb* are a consequence of presynaptic Ca^2+^ dyshomeostasis due to the inability of the neurons to maintain the levels of ATP needed for ensuring Ca^2+^ extrusion. Our findings agree with the insights gleaned from *in silico* modeling of ALS neurons (Le Masson et al., 2014), and provide an overarching explanation for the presynaptic defects associated with the expression of the ALS-causing variant of *vapb* in *Drosophila* motor neurons.

## Results

### Expression of an ALS-related variant of *vapb* results in defective presynaptic bouton development

Either the deletion of *Drosophila vapb* (also called *Vap33*) or the expression of a transgene equivalent to an ALS-causing variant (*vapb^P58S^*) disrupts presynaptic microtubules at the larval NMJ resulting in the appearance of fewer, but morphologically larger, boutons (Pennetta et al., 2002; Chai et al., 2008; Ratnaparkhi et al., 2008). We confirmed that ectopic expression of *vapb^P58S^*, but not wild-type *vapb* (*vapb^WT^*), in motor neurons (using *VGlut^ok371^-GAL4*, herein referred to as *ok371-GAL4*) led to a significant reduction in bouton number (Fig. 1A-E) (Brand and Perrimon, 1993; Mahr and Aberle, 2006; Tsuda et al., 2008). NMJ boutons in neurons expressing *vapb^P58S^* also exhibited significant increases in bouton area (Fig. 1A-B and 1F). Direct comparison with animals expressing *vapb^WT^* revealed that those expressing *vapb^P58S^* exhibited significantly fewer and larger boutons (Fig. 1E-F). These data agree with prior findings regarding the effects of VAPB^P58S^ on *Drosophila* larval NMJ synapse morphology (Chai et al., 2008; Ratnaparkhi et al., 2008).

**FIGURE 1.**
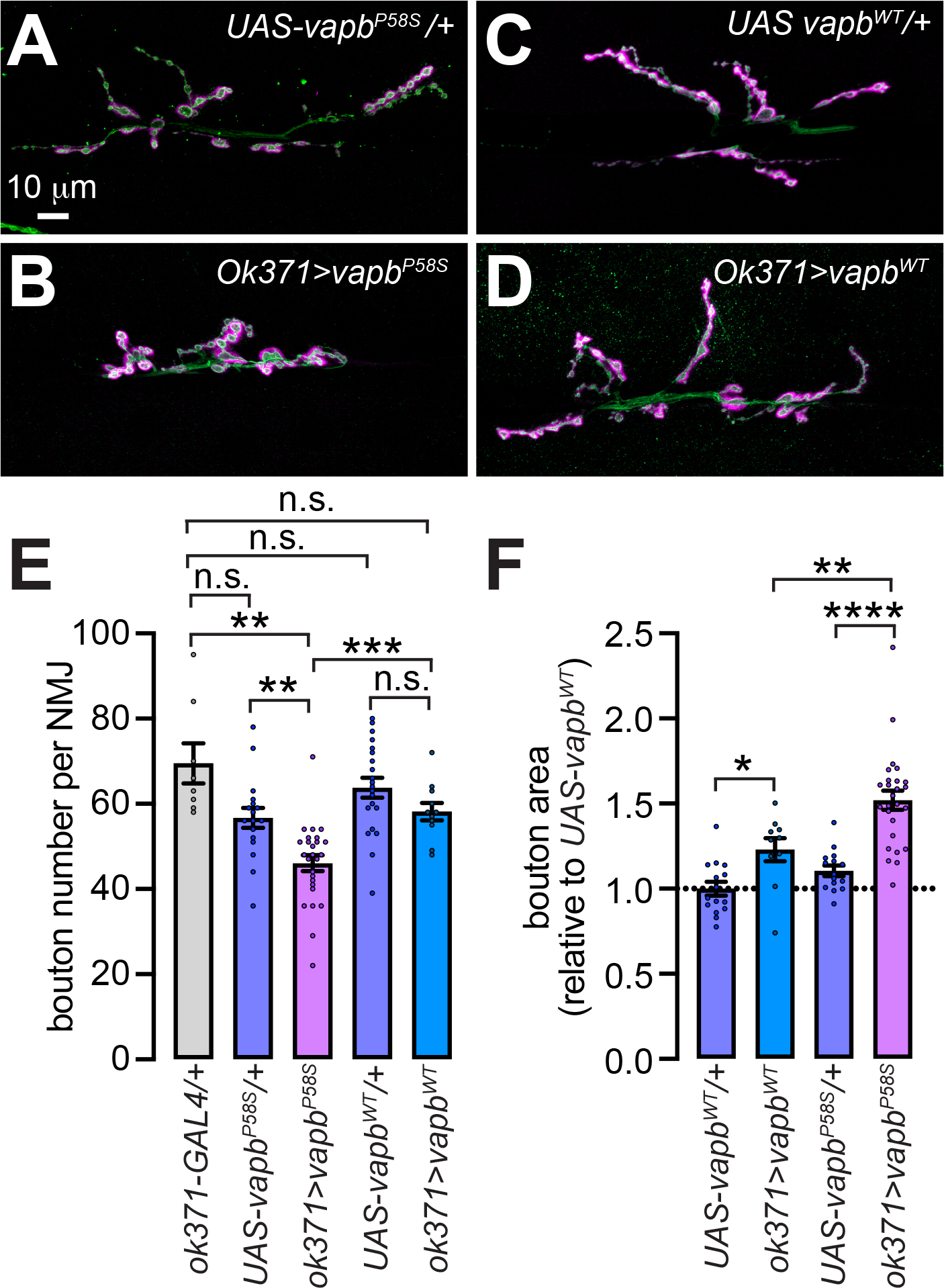
Expression of *vapb^P58S^* in *Drosophila* motor neurons led to significant changes in presynaptic bouton development at the larval NMJ. **(A-D)** Representative confocal images of larval NMJs dissected from animals of the indicated genotypes stained with antibodies against HRP (green) and DLG (magenta). Scale bar shown in **(A)** on the top left applies to all panels. Please note that in all figures, “*UAS-transgene*/+” refer to the presence of the noted *UAS-transgene* construct without a *GAL4* driver, and “*driver>transgene*” refers to the *UAS-transgene*, whose expression was driven using the *driver-GAL4*. **(E-F)** Bar graphs showing quantification of the larval NMJ bouton numbers **(E)**and relative bouton area **(F)** in animals of the indicated genotypes. Values represent mean ± SEM. **, *P*<0.01; ***, *P*<0.001; ****, *P*<0.0001; n.s., not significant, t-tests with Bonferroni correction. Dots represent values from distinct NMJs.

### VAPB^P58S^-induced defects in presynaptic bouton development are ameliorated by lowering the expression of genes encoding ER Ca^2+^ release channels

We previously showed that decreased abundance of ER Ca^2+^ release channels — Inactive (Iav), ryanodine receptor (RyR), and inositol trisphosphate receptor (IP_3_R) — result in NMJs that exhibit fewer boutons that are larger in size (Fig. 2A) (Wong et al., 2014). This implied ostensible similarities between the morphology of NMJ boutons in *iav^1^* mutants (also called *iav^hypoB-1^* — strong hypomorphs of *iav* (Homyk and Sheppard, 1977; Wong et al., 2014)) and animals expressing *vapb^P58S^*. Therefore, we speculated that the coincidence of *iav^1^* and *vapb^P58S^* would enhance the respective phenotypes leading to an even greater reduction in bouton number (Fig. 2A). To our surprise, expression of *vapb^P58S^* in the *iav^1^* background restored, rather than worsened, both bouton number and morphology to control levels (Fig. 2B-C and 2J-K). Whereas bouton numbers and area elicited by *vapb^P58S^* were not significantly different from those in *iav^1^*, these parameters were restored to wild-type levels upon the coincidence of *vapb^P58S^* and *iav^1^* (Fig. 2J-K). In contrast, overexpression of *vapb^WT^* did nothing to alter the paucity of boutons in *iav^1^* (Fig. 2D-E and 2L). Therefore, decreased abundance of Iav rescued the synaptic growth phenotype stemming from the expression of *vapb^P58S^*.

**FIGURE 2.**
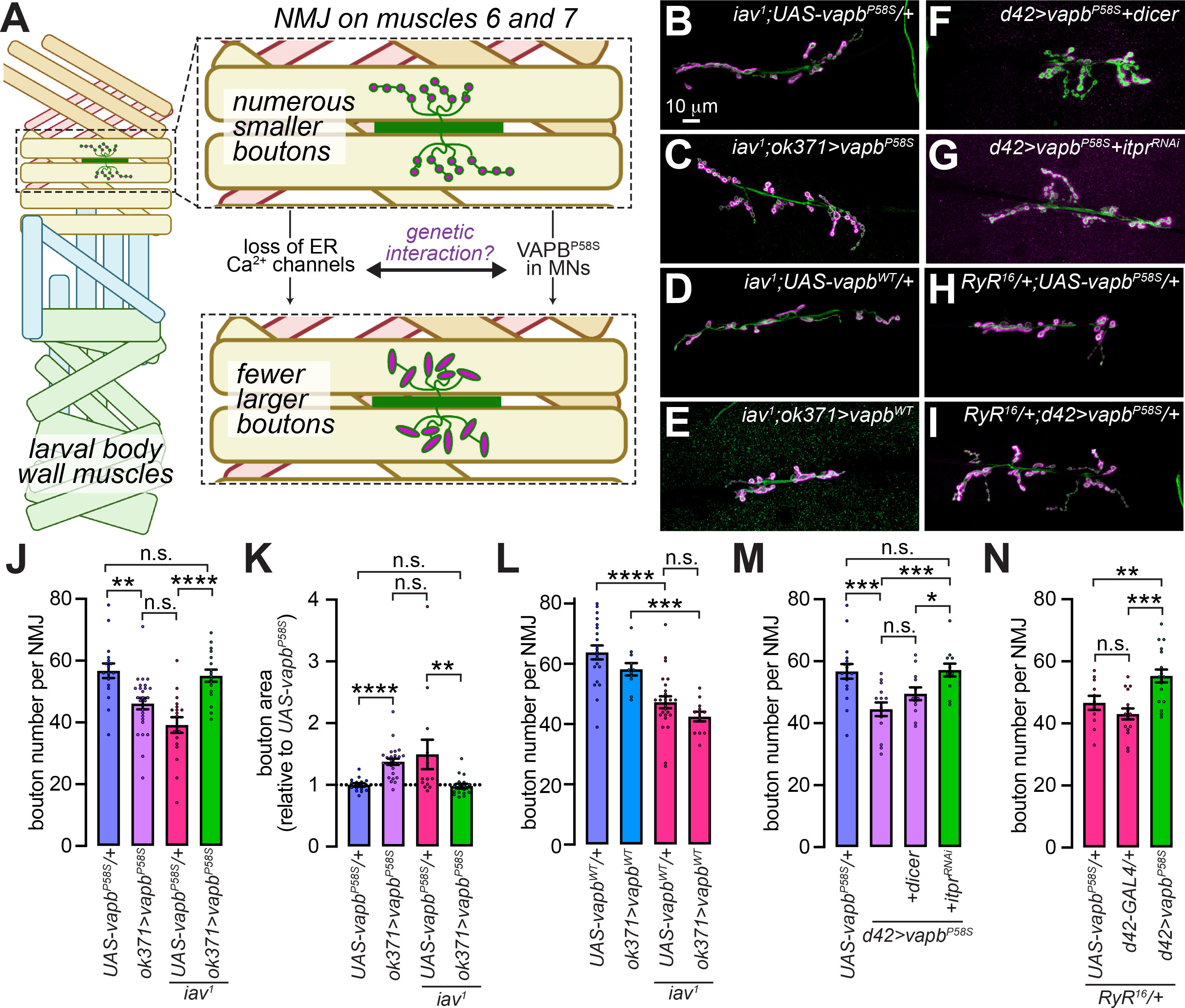
VAPB^P58S^-induced defects in presynaptic bouton development are ameliorated upon concomitant loss of ER Ca^2+^ release channels. **(A)** Model showing that the development of control NMJs on muscles 6 and 7 of the larval body wall muscles leads to the appearance of numerous small boutons. Animals either expressing *vapb^P58S^* in motor neurons or harboring mutations that diminish ER Ca^2+^ release are characterized by the appearance of fewer, but morphologically larger, boutons. Ostensible similarities in bouton development phenotypes raises the question of genetic interactions between these conditions. Image was created with BioRender.com. **(B-I)** Representative confocal images of larval NMJs dissected from animals of the indicated genotypes stained with antibodies against HRP (green) and DLG (magenta). Scale bar shown in **(B)** applies to all panels. **(J-N)** Bar graphs showing quantification of the larval NMJ bouton numbers **(J, L-N)**and relative bouton area **(K)** in animals of the indicated genotypes. Values represent mean ± SEM. *, *P*<0.05; **, *P*<0.01; ***, *P*<0.001; ****, *P*<0.0001; n.s., not significant, paired t-tests with Bonferroni correction. Dots represent values from distinct NMJs.

Next, we asked whether the rescue of *vapb^P58S^*-induced NMJ phenotypes was specific to the loss of Iav, or whether similar suppression would occur in response to lowered abundance of other ER Ca^2+^ channels. Simultaneous knockdown of the gene encoding the fly IP_3_R (*itpr*) using the *d42-GAL4* motor neuron driver and a validated RNAi line (*UAS-itpr^RNAi^*) ameliorated the effect of *vapb^P58S^* expression on bouton number (Fig. 2G and 2M) (Venkatesh and Hasan, 1997; Parkes et al., 1998; Wong et al., 2014, 2021). Absence of comparable suppression in animals coexpressing *vapb^P58S^* and a neutral *UAS* transgene (*UAS-Dcr1*) (Fig. 2F and 2M) demonstrates that the suppression brought about by coexpression of *itpr^RNAi^* was not due to GAL4 dilution stemming from the presence of a second *UAS* transgene. Concomitant absence of one copy of *RyR* (*RyR^16^*/+), which mimics the *iav^1^* NMJ growth phenotype, also prevented *vapb^P58S^*-induced alterations in bouton numbers (Fig. 2H-I and 2N) (Sullivan et al., 2000; Wong et al., 2014). Taken together, these data demonstrate that VAPB^P58S^-induced defects in presynaptic bouton development are ameliorated by cell autonomous reduction in the expression of genes encoding ER Ca^2+^ channels.

### CaMKII overactivation underlies alterations in presynaptic development in motor neurons expressing *vapb^P58S^*

The aforementioned findings are consistent with the notion of VAPB^P58S^ inducing an increase in cytosolic [Ca^2+^] such that the attenuation of ER Ca^2+^ release restores homeostasis, and thus, mitigates the bouton development phenotypes. Our data also imply that either an increase or decrease in presynaptic [Ca^2+^] results in morphologically identical bouton phenotypes. Indeed, we previously showed that either the absence or overexpression of *iav* results in comparable reduction in the number of NMJ boutons (Wong et al., 2014).

### How might this bell-shaped relationship between presynaptic [Ca^2+^] and NMJ bouton development come about?

We posit that the key to understanding these outcomes is the phosphorylation status of the *Drosophila* microtubule-associated protein-1b homolog, Futsch — an important determinant of presynaptic microtubule stability and NMJ bouton morphology (Hummel et al., 2000; Wong et al., 2014). While dephosphorylated Futsch binds and stabilizes presynaptic microtubules, phosphorylated Futsch dissociates from microtubules leading to depolymerization of those cytoskeletal filaments (Fig. 3A). Results from many groups are consistent with either the hyperphosphorylation or absence of Futsch leading to a loss of presynaptic microtubules, increased bouton size, and decreased bouton number (Roos et al., 2000; Pennetta et al., 2002; Gögel et al., 2006; Viquez et al., 2006; Wong et al., 2014). Of relevance to the present study, the phosphorylation status of Futsch appears to depend on cytosolic [Ca^2+^]. Reduction in presynaptic resting [Ca^2+^] and the attendant attenuation of the Ca^2+^/calmodulin (CaM) responsive phosphatase, calcineurin (CanA1), destabilize presynaptic microtubules in a Futsch-dependent manner (Wong et al., 2014). In principle, increased activity of a Ca^2+^-responsive kinase could also result in hyperphosphorylated Futsch. If so, one may expect that either an increase or decrease in cytosolic [Ca^2+^] would result in higher fractions of phosphorylated Futsch, and thus, microtubule depolymerization (Fig. 3A).

**FIGURE 3.**
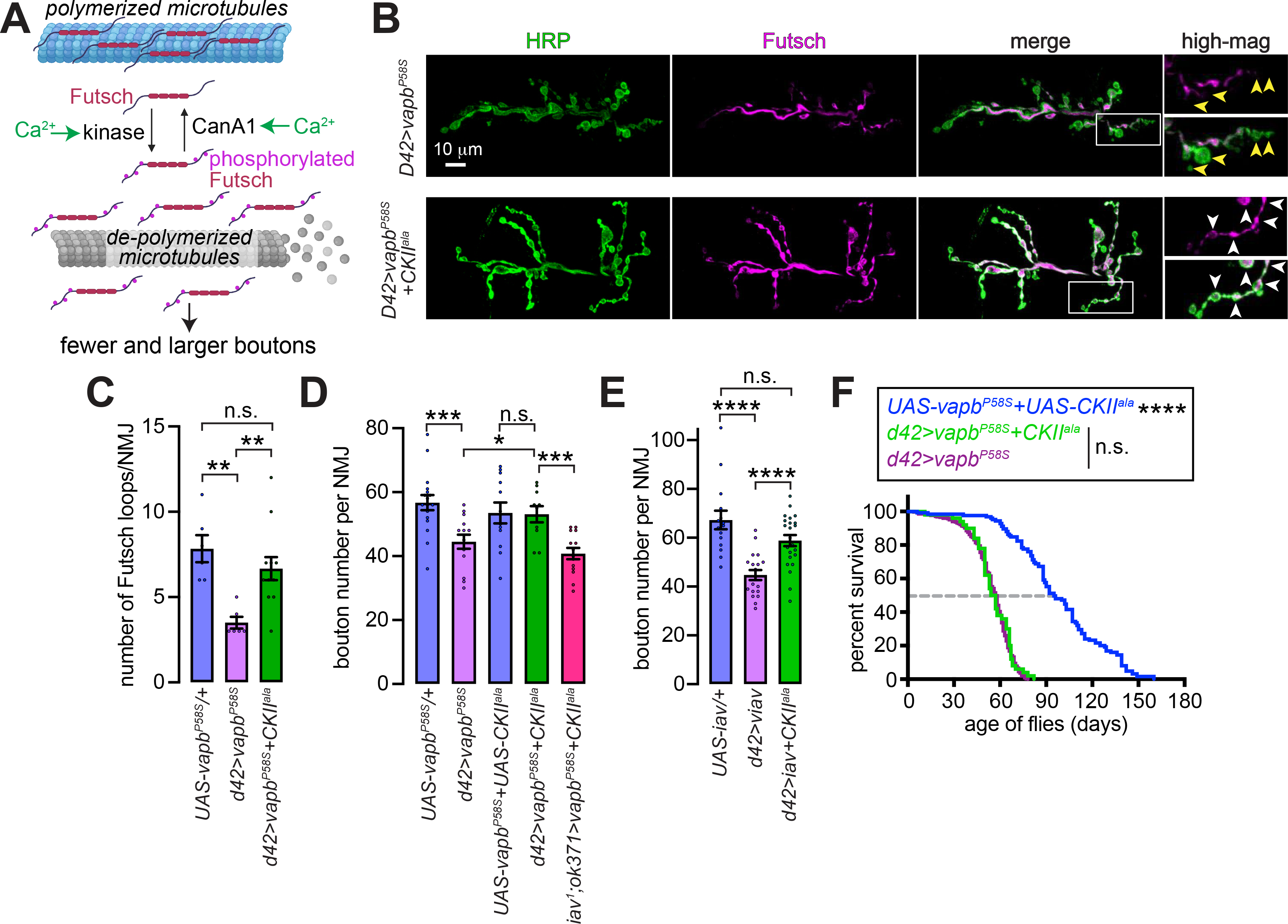
CaMKII underlies the alterations in presynaptic bouton development in motor neurons expressing *vapb^P58S^*. **(A)** Model showing that either an increase or decrease in presynaptic [Ca^2+^] can result in Futsch hyperphosphorylation and depolymerization of presynaptic microtubules. Elevated [Ca^2+^] would lead to persistent activation of CaMKII, which could induce Futsch phosphorylation and attendant disruption of presynaptic microtubules leading to the appearance of fewer, but larger, boutons. Decreased expression of ER Ca^2+^ channels results in lower presynaptic resting [Ca^2+^] and diminished calcineurin activity (Wong et al., 2014), which also results in Futsch phosphorylation and attendant disruption of presynaptic microtubules leading to the appearance of fewer, but larger, boutons. Image was created with BioRender.com. **(B)** Representative confocal images of NMJs expressing *vapb^P58S^* either with or without *KII^ala^* as indicated. Samples were stained with antibodies against HRP (green) and Futsch (magenta). White arrowheads point to individual boutons and highlight recovery of Futsch loops in neurons expressing both *vapb^P58S^* and *CKII^ala^*. Higher magnification images (high-mag) represent the regions shown in the box overlaid on the merged images. Scale bar shown on the top left applies to all panels. **(C-E)** Bar graphs showing quantification of the number of Futsch loops per NMJ **(C)** and larval NMJ bouton numbers **(D-E)** in animals of the indicated genotypes. Values represent mean ± SEM. *, *P*<0.05; **, *P*<0.01; ***, *P*<0.001; ****, *P*<0.0001; n.s., not significant, paired t-tests with Bonferroni correction. Dots represent values from distinct NMJs. **(F)** Lifespan of flies of the indicated genotypes. ****, *P*<0.0001; n.s., not significant, log- rank tests with Bonferroni correction.

Putative increase in Futsch phosphorylation has been linked to the loss of Futsch loops within presynaptic boutons (Viquez et al., 2006; Wong et al., 2014). Motor neurons expressing *vapb^P58S^* also exhibited fewer Futsch loops within presynaptic boutons (Fig. 3B-C). Since activation of the Ca^2+^/CaM-dependent protein kinase II (CaMKII or CKII) by high [Ca^2+^] destabilizes microtubules by phosphorylating microtubule-associated proteins (Baratier et al., 2006; McVicker et al., 2015; Oka et al., 2017), we asked whether elevated ER Ca^2+^ release compels the changes in NMJ development due to unrestrained CaMKII activity. Coexpression of the CaMKII inhibitory peptide, CaMKII^ala^ (CKII^ala^) (Joiner MlA and Griffith, 1997), with *vapb^P58S^* led to recovery of the numbers of Futsch loops (Fig. 3B-C) and boutons (Fig. 3D). In *iav^1^* synapses, which exhibit lower resting [Ca^2+^] (Wong et al., 2014), expression of *CKII^ala^* did not restore the bouton number (Fig. 3D). Indicating the sufficiency of CaMKII downstream of elevated [Ca^2+^], in *iav*-overexpressing neurons, which exhibit higher presynaptic [Ca^2+^] (Wong et al., 2014), bouton number was restored to wild-type level upon coexpression of *CKII^ala^* (Fig. 3E). Taken together, these data suggest that *vapb^P58S^*-induced alterations in presynaptic development stem from aberrant CaMKII activation.

In a recent study, we showed that expression of *vapb^P58S^* in motor neurons shortens *Drosophila* lifespan via overactivation of PLC®–IP_3_R signaling (Wong et al., 2021). Consequently, either the deletion of the gene encoding a fly PLC® (*norpA*) or the concomitant knockdown of *itpr* in neurons expressing *vapb^P58S^* strongly suppressed the effects of VAPB^P58S^ on animal longevity (Bloomquist et al., 1988; Wong et al., 2021). In contrast to the ameliorative effects on the NMJ bouton phenotype, neither *iav^1^* nor the *RyR^16^*/+ mutations influenced the lifespan of animals expressing *vapb^P58S^* (Wong et al., 2021). Therefore, the effects of *vapb^P58S^* on NMJ bouton development and adult lifespan occur via distinct mechanisms. In agreement, coexpression of *CKII^ala^*, which mitigated the NMJ growth phenotype in animals expressing *vapb^P58S^*, did not influence the effect of mutant VAPB on adult lifespan (Fig. 3F).

### Expression of *vapb^P58S^* results in diminished extrusion of cytosolic Ca^2+^

Lower presynaptic resting [Ca^2+^] in the absence of Iav results in reduced probability of synaptic vesicle (SV) release (Wong et al., 2014). NMJs in *iav^1^* animals, therefore, exhibit reduced amplitudes of evoked excitatory postsynaptic junctional potentials (EJPs) and paired-pulse facilitation in response to a 2^nd^ stimulus applied after a delay of 50 milliseconds (Wong et al., 2014). Conversely, overexpression of *iav* led to an increase in presynaptic resting [Ca^2+^], higher EJP amplitudes, and paired-pulse depression indicating elevated SV release probability (Wong et al., 2014). These data point to a dose-dependent effect of ER Ca^2+^ release on presynaptic resting [Ca^2+^] and SV release probability. Given that the NMJ bouton phenotypes in animals overexpressing either *vapb^P58S^* or *iav* were suppressed by CKII^ala^, we asked whether VAPB^P58S^ also elevates SV release probability due to an increase in resting [Ca^2+^]. If so, expression of *vapb^P58S^* would result in increased EJP amplitude and paired-pulse depression. However, EJP amplitudes in *vapb^P58S^* neurons were not significantly different from those in *vapb^WT^* neurons (Fig. 4A-B), which is in agreement with the report that ectopic expression of the ALS-causing human transgene (*hVAP^P56S^*) had no effect on the amplitude of evoked potentials at the *Drosophila* larval NMJ (Chai et al., 2008).

**FIGURE 4.**
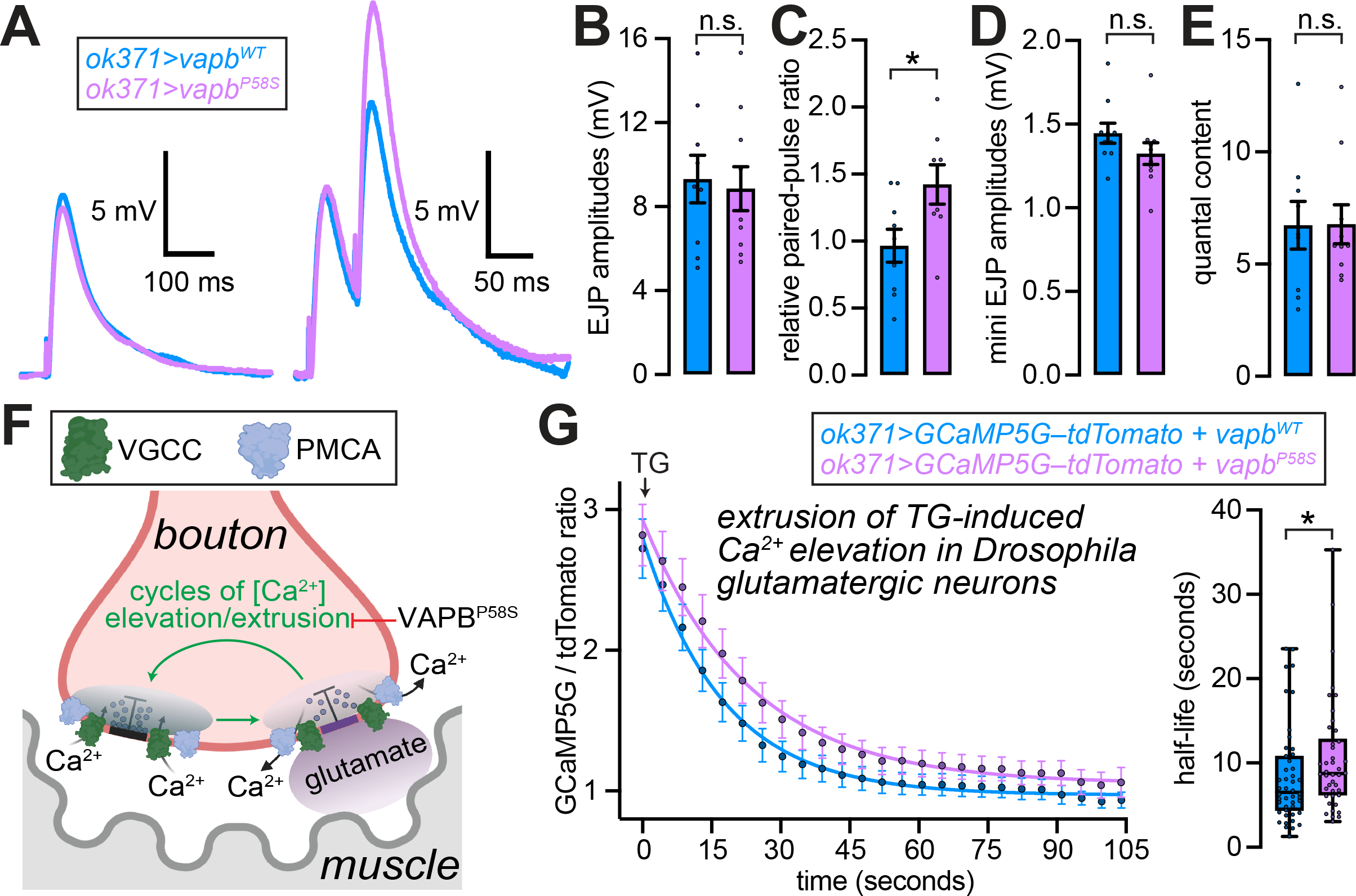
Expression of *vapb^P58S^* results in delayed extrusion of cytosolic Ca^2+^. **(A)** Representative EJP (left) and paired-pulse EJP (right) traces recorded from larval NMJs isolated from animals expressing *vapb^WT^* (blue trace) or *vapb^P58S^* (magenta trace) in motor neurons using *ok731-GAL4*. **(B)** Bar graph showing quantification of the EJP amplitudes from the data shown in **(A)**. Values represent mean ± SEM. n.s., not significant, paired t-tests. Dots represent values from distinct NMJs. **(C)** Bar graph showing quantification of the paired-pulse ratio (fractional change in the amplitude of the second EJP to that of the first EJP when the two stimulatory pulses were applied 50ms apart) of the data shown in **(A)**. Values represent mean ± SEM. *, *P*<0.05, t-tests. Dots represent values from distinct NMJs. **(D)** Bar graph showing quantification of the mini EJP amplitudes. Values represent mean ± SEM. n.s., not significant, paired t-tests. Dots represent values from distinct NMJs. **(E)** Bar graph showing quantification of quantal content (ratio of amplitudes of EJP and mini EJP). Values represent mean ± SEM. n.s., not significant, paired t-tests. Dots represent values from distinct NMJs. **(F)** Model showing that the rates of [Ca^2+^] elevation and extrusion are necessary for maintaining the fidelity of synaptic transmission. Paired-pulse facilitation without a change in the amplitude of the first pulse, as seen in **(A)**, could be explained by VAPB^P58S^ decreasing the rates of Ca^2+^ extrusion. Image was created with BioRender.com. **(G)** *Left*, traces showing the decay of GCaMP5G/tdTomato ratio after thapsigargin (TG)- induced cytosolic [Ca^2+^] elevations in motor neurons dissociated from animals of the indicated genotypes. Values were fit to a 1^st^ order exponential function. Values represent mean ± SEM of traces from >50 neurons of each genotype. *Right*, box plots showing quantification of half-lives of decay of the signals in individual neurons of the indicated genotypes. *, *P*<0.05, t-tests, Mann-Whitney test. Dots represent values from individual neurons expressing the indicated transgenes.

Furthermore, relative to neurons expressing *vapb^WT^*, those expressing *vapb^P58S^* showed paired-pulse facilitation, rather than paired-pulse depression (Fig. 4A and 4C).

Amplitudes of mini-EJPs (mEJPs) and quantal content were also virtually identical in *vapb^WT^*- and *vapb^P58S^*-expressing NMJs (Fig. 4D-E). Taken together, these data argue against elevations in presynaptic resting [Ca^2+^] and SV release probability in *vapb^P58S^*- expressing motor neurons.

### What then could explain the hallmarks of elevated [Ca^2+^] in vapb^P58S^-expressing motor neurons

Elevations in cytosolic [Ca^2+^] could stem either from greater Ca^2+^ influx or from diminished Ca^2+^ extrusion from the cytosol. Since the former is ruled out by the aforementioned data, we asked whether the effects of VAPB^P58S^ stem from diminished Ca^2+^ extrusion (Fig. 4F). We reasoned that diminished Ca^2+^ extrusion would promote the accumulation of residual Ca^2+^, boost SV release in response to rapidly delivered stimuli, and thereby, explain the paired-pulse facilitation in *vapb^P58S^*-expressing neurons (Fig. 4A and 4C). To examine the rates of Ca^2+^ extrusion, we performed live-cell imaging of larval motor neurons expressing GCaMP5G-tdTomato (Daniels et al., 2014; Wong et al., 2014, 2021). To ensure that our assessment of Ca^2+^ extrusion focuses on transfer to the extracellular medium rather than uptake into the ER, we applied the SERCA inhibitor, thapsigargin (TG) (Lytton et al., 1991; Wong et al., 2021). Incidentally, TG-treatment evokes cytosolic Ca^2+^ transients due to leak of ER stores into the cytosol (Wong et al., 2021), and the kinetics of the return of these transients to baseline reflect the rate of Ca^2+^ extrusion. We found that the decay of TG-evoked GCaMP5G-tdTomato elevations followed a 1^st^ order exponential function (Fig. 4G). Half-life of this decay was significantly higher in neurons expressing *vapb^P58S^* compared to those expressing *vapb^WT^* (Fig. 4G). These data argue in favor of diminished Ca^2+^ extrusion in neurons expressing *vapb^P58S^*.

### Rates of ATP production are unable to keep up with demand in *vapb^P58S^*- expressing motor neurons

Electrochemical homeostasis in healthy neurons is maintained by pumps powered by mitochondrially-derived ATP. While SERCA and PMCA restore depolarization-induced changes in cytosolic [Ca^2+^] to resting levels, the Na^+^/K^+^ ATPase restores the membrane potential (Fergestad et al., 2006; Chouhan et al., 2012; Ivannikov and Macleod, 2013; Le Masson et al., 2014). The purported coupling of ATP production to neuronal activity, therefore, ensures the availability of ATP for maintenance of Ca^2+^ homeostasis and for membrane repolarization (Fig. 5A, left) (Le Masson et al., 2014; Rangaraju et al., 2014; Wong et al., 2021). In ALS neurons, defects in mitochondrial function are predicted to result in a paucity of ATP, which in turn, attenuates the activity of Ca^2+^- and Na^+^/K^+^ ATPases (Fig. 5A, right) (Le Masson et al., 2014). *In silico* modeling has revealed that the outcome of mitochondrial dysfunction and diminished pump activity is a “deadly loop” comprised of a rapidly burgeoning demand for ATP that remains unmet, and Ca^2+^ dyshomeostasis (Fig. 5A, right) (Le Masson et al., 2014). Since our findings of diminished Ca^2+^ extrusion in neurons expressing *vapb^P58S^* (Fig. 4G) align with the notion of Ca^2+^ dyshomeostasis, we asked whether the expression of *vapb^P58S^* — ***(1)*** compromises mitochondrial ATP production, and thereby, precludes a suitable response to the ATP burden of depolarization; and ***(2)*** increases the rates of overall ATP hydrolysis as inferred from *in silico* modeling of ALS neurons (Le Masson et al., 2014).

**FIGURE 5.**
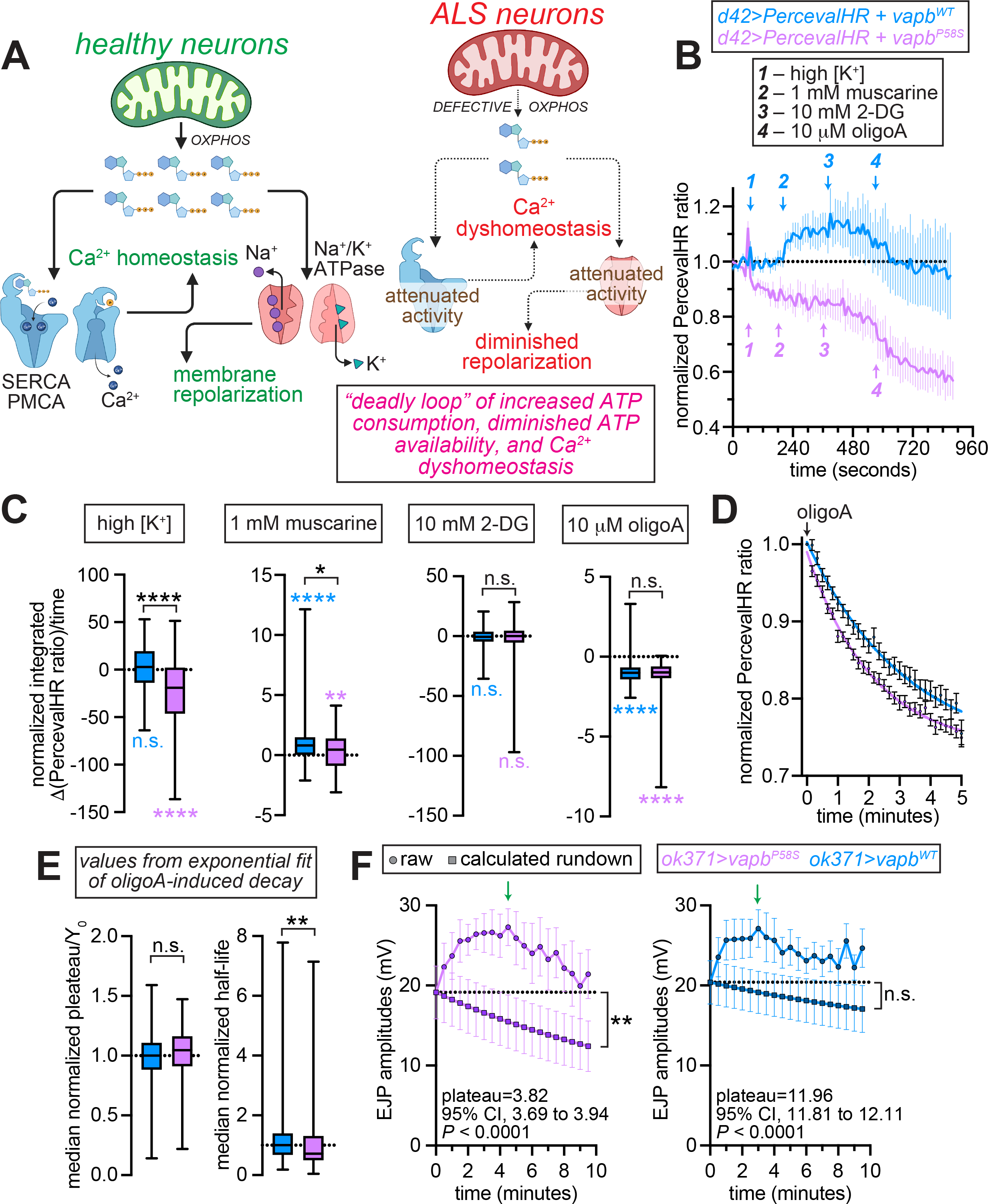
Activity-dependent ATP production and synaptic transmission is diminished in motor neurons expressing *vapb^P58S^*. **(A)** Model adapted from Le Masson et al. (Le Masson et al., 2014) showing the role for neuronal ATP in mediating ionic homeostasis. In healthy neurons, ATP derived from mitochondrial oxidative phosphorylation (OXPHOS) powers Ca^2+^ ATPases, such as SERCA and PMCA, to maintain Ca^2+^ homeostasis, and the Na^+^/K^+^ ATPase, which is needed for setting the resting membrane potential and for repolarization of the membrane potential after bouts of depolarization. In ALS neurons, a decrease in mitochondrial ATP production would attenuate the activities of Ca^2+^- and Na^+^/K^+^- ATPases. This would be predicted to set in motion a self-reinforcing “deadly loop” comprised of continuously increasing ATP consumption coupled with diminished ATP availability, Ca^2+^ dyshomeostasis, and the eventual loss of membrane potential. Image was created with BioRender.com. **(B)** Representative traces showing normalized PercevalHR ratio in *Drosophila* motor neurons coexpressing the indicated transgenes. Values represent mean ± SEM of traces from 14-15 neurons of each genotype. Arrows indicate treatments. **(C)** Box plots showing the integrated change in the PercevalHR ratio per unit time. Median value of zero indicates no change in the PercevalHR ratio (i.e., [ATP]/[ADP] ratio), median values > 0 denote a net increase in the PercevalHR ratio after the treatment, and median values < 0 denote a net decrease in the PercevalHR ratio after the treatment. *, *P*<0.05; **, *P*<0.01; ***, *P*<0.001; ****, *P*<0.0001; n.s., not significant. Colored significance values below the box plots show results of one-sample Wilcoxon test for the hypothetical median value of 0. Significance tests shown on the top represent results of Mann-Whitney tests for comparing median values in the two genotypes. **(D)** Traces showing the decay of the PercevalHR ratio after application of oligomycin A (oligoA). The time of oligoA addition was set to 0. Values were normalized to the PercevalHR ratio at the time of oligoA addition, and represent mean ± SEM of traces from 85-120 neurons of each genotype. Values were also fit to 1^st^ order exponential functions. **(E)** Parameters of the fit to 1^st^ order exponential functions from **(D)**. *Left*, box plot showing the normalized ratio of the plateau and starting (Y_0_) values. *Right*, box plot showing the normalized half-life of decay. **, *P*<0.01 and n.s., not significant, Man-Whitney tests. **(F)** Traces showing EJP amplitudes recorded from NMJs of animals of the indicated genotypes stimulated at 10 Hz for 10 minutes. Traces connecting filled circles represent raw values that were determined experimentally, and the traces connecting filled squares represent calculated rundowns fit to 1^st^-order exponential decay. Green arrows indicate the onset of decay. *CI*, confidence interval; *P*-values for CI were calculated as described in the methods. All values represent mean ± SEM of EJP values recorded from 5 NMJ preparations of each genotype.

Given that mitochondrial Ca^2+^ uptake — needed for energizing those organelles and stimulating TCA dehydrogenases — is disrupted in *vapb^P58S^*-expressing neurons, it stood to reason that “on-demand” ATP production would be attenuated in these neurons (Duchen, 1992; McCormack and Denton, 1993; Dumollard et al., 2004; Cárdenas et al., 2010; Ding et al., 2018; Wong et al., 2021). To directly assess ATP production and consumption in neurons, we monitored the cytosolic [ATP]/[ADP] ratio using the genetically-encoded sensor, PercevalHR (Tantama et al., 2013; Wong et al., 2021). In neurons coexpressing *vapb^WT^*, the PercevalHR ratio (i.e., the [ATP]/[ADP] ratio) remained unchanged in response to depolarization by the application of high [K^+^] (52 mM) (Fig. 5B, ***blue trace – 1***). These data, which agree with our prior findings in wild-type neurons (Wong et al., 2021), indicate that the rates of ATP production and consumption were perfectly balanced in *vapb^WT^*-expressing neurons following depolarization. In *vapb^P58S^*-expressing neurons, however, high [K^+^] led to a decline in the PercevalHR ratio (Fig. 5B, ***magenta trace – 1***). Comparison of the integrated change in the PercevalHR ratio per unit time revealed a highly significant decrease from baseline in *vapb^P58S^*-expressing neurons, but a statistically insignificant change in *vapb^WT^* neurons (Fig. 5C, ***high [K^+^]***). Furthermore, the integrated drop in the PercevalHR ratio in *vapb^P58S^* neurons was significantly greater than that in *vapb^WT^* neurons (Fig. 5C, ***high [K^+^]***). These data indicate that *vapb^P58S^*-expressing neurons were unable to maintain the [ATP]/[ADP] ratio after depolarization, which pointed to enhanced ATP consumption and/or compromised response to the ATP burden of depolarization.

We previously showed that in *vapb^WT^*-expressing neurons, IP_3_R activation in response to muscarine-induced stimulation of PLC®-coupled acetylcholine receptors elevates mitochondrial [Ca^2+^] as a consequence of interorganellar Ca^2+^ transfer (Wong et al., 2021). Consistent with reports of ALS-causing variants of VAPB disrupting interorganellar Ca^2+^ transfer, muscarine-evoked mitochondrial [Ca^2+^] elevations were absent in *vapb^P58S^*-expressing neurons (De Vos et al., 2012; Stoica et al., 2016; Gomez-Suaga et al., 2017; Smith et al., 2017; Xu et al., 2020; Wong et al., 2021). Given the role for matrix [Ca^2+^] in energizing mitochondria and promoting ATP production (Duchen, 1992; McCormack and Denton, 1993; Dumollard et al., 2004; Cárdenas et al., 2010; Ding et al., 2018), we reasoned that diminished interorganellar Ca^2+^ transfer would attenuate muscarine-induced ATP production in *vapb^P58S^* neurons. Indeed, muscarine evoked a significantly smaller elevation in the [ATP]/[ADP] ratio in *vapb^P58S^* neurons than it did in neurons expressing *vapb^WT^* (Fig. 5B, ***2*** and 5C, ***1 mM muscarine***). These data support the principle that diminished Ca^2+^ uptake into the mitochondrial matrix perturbs the rate of ATP production in *vapb^P58S^*-expressing neurons.

Subsequent inhibition of glycolysis by the application of 2-deoxyglucose (2-DG) did not affect the [ATP]/[ADP] ratio in neurons expressing either *vapb^WT^* or *vapb^P58S^* (Fig. 5B, ***3*** and 5C, ***10 mM 2-DG***). In contrast, inhibition of OXPHOS using the ATP synthase inhibitor, oligomycin A (oligoA), decreased the [ATP]/[ADP] ratio in neurons expressing either *vapb^WT^* or *vapb^P58S^* (Fig. 5B, ***4*** and 5C, ***10*** *μ**M oligoA***). Therefore, the [ATP]/[ADP] ratio in those neurons exhibited greater sensitivity to the inhibition of OXPHOS rather than glycolysis. The drops in the integrated PercevalHR ratio after inhibition of OXPHOS were comparable in the *vapb^WT^* and *vapb^P58S^* neurons (Fig. 5C, ***10*** *μ**M oligoA***). These data imply that upon the cessation of ATP production, the cumulative drop in the [ATP]/[ADP] ratio is the same in neurons expressing either *vapb^WT^* or *vapb^P58S^*. Therefore, the decline in the [ATP]/[ADP] ratio in depolarized *vapb^P58S^* neurons reflects changes in the rates of ATP synthesis and consumption rather than changes in ADP availability. In agreement, upon fitting the decay of the [ATP]/[ADP] ratio to 1^st^ order exponential functions (Fig. 5D), we found that the [ATP]/[ADP] ratio settles at comparable values in *vapb^WT^* and *vapb^P58S^* neurons (i.e., insignificant differences in plateau/Y_0_ values) (Fig. 5E). However, the rate of decay was significantly greater in *vapb^P58S^*, resulting in a shorter half-life (Fig. 5E). These data imply elevated rates of ATP consumption in *vapb^P58S^*-expressing *Drosophila* motor neurons.

### Loss of synaptic transmission during high-frequency stimulation in *vapb^P58S^*- expressing motor neurons

Many *vapb^P58S^* phenotypes we describe in this study, including elevated cytosolic [Ca^2+^] and ATP depletion, are also observed in *drp1*-deficient animals (Verstreken et al., 2005). Since the paucity of ATP in *drp1* mutant NMJs precludes the maintenance of synaptic transmission during high-frequency stimulation (Verstreken et al., 2005), we asked whether the shortage of ATP in *vapb^P58S^* motor neurons would result in similar rundown of SV release. Although stimulation of the *vapb^P58S^* NMJs at 10 Hz led to EJP amplitudes that initially plateaued at values that were ∼35% *higher* than baseline, those values transitioned to periods of sustained decay after ∼5 minutes of high-frequency stimulation (Fig. 5F left, traces with filled circles, green arrow indicates onset of decay). EJP amplitudes in 100% of the traces recorded from *vapb^P58S^* animals stimulated at 10 Hz exhibited sustained periods of decay after initial, transient elevations. We reasoned that superimposed on the liminal increase in EJP amplitudes were the rundowns of SV release, which become apparent after ∼5 minutes of high-frequency stimulation. We extracted the rundown components from the traces using the rate-constants of the decay (Fig. 5F left, traces with filled squares). Over the 10-minute duration of the high-frequency stimulation, calculated rundowns in *vapb^P58S^* NMJs dropped significantly lower than baseline (Fig. 5F, left). The decay phase of high-frequency traces recorded from *vapb^WT^* NMJs was relatively less-pronounced (Fig. 5F, right), with only 40% of the traces exhibiting sustained periods of declining EJP values. Calculated rundown in *vapb^WT^* NMJs was not significantly lower than baseline (Fig. 5F, right). Furthermore, calculated rundowns in *vapb^P58S^* were predicted to plateau at 3.82 mV (95% CI, 3.69 to 3.94, *P* < 0.0001), whereas rundowns were predicted to plateau at 11.96 mV (95% CI, 11.81 to 12.11, *P* < 0.0001) in *vapb^WT^*. Taken together, these data indicate that high-frequency stimulation leads to a greater loss of synaptic transmission in *vapb^P58S^*- expressing motor neurons.

## Discussion

Our findings explain the characteristic changes to NMJ morphology observed in animals that harbor *vapb* deletions or express an ALS-causing variant (*vapb^P58S^*) of the gene (Pennetta et al., 2002; Chai et al., 2008; Ratnaparkhi et al., 2008). While it was known that the reduction in NMJ bouton number and the increase in bouton size in these animals stemmed from diminished stability of the presynaptic microtubules (Pennetta et al., 2002; Chai et al., 2008; Ratnaparkhi et al., 2008), the molecular underpinnings of these cytoskeletal changes had remained incompletely understood. Our findings that the morphological defects observed in *vapb^P58S^* NMJs were attenuated by the coexpression of a peptide that inhibits CaMKII implicate the aberrant activation of this kinase in the appearance of these phenotypes. These conclusions agree with prior reports of CaMKII regulating microtubule stability by phosphorylating microtubule-associated proteins (Baratier et al., 2006; McVicker et al., 2015; Oka et al., 2017).

We also present data that raise the possibility that the MAP1b homolog, Futsch (Hummel et al., 2000; Roos et al., 2000), could be the relevant CaMKII target in this context. Previous studies have shown that hyperphosphorylation of Futsch, which provokes the destabilization of presynaptic microtubules, results in the loss of Futsch- loops within the NMJ boutons (Gögel et al., 2006; Viquez et al., 2006; Wong et al., 2014). Not only did the *vapb^P58S^* NMJs exhibit fewer Futsch loops, these values were restored to wild-type levels upon the coexpression of the CaMKII inhibitory peptide. The involvement of Futsch in the phenotypes observed in this model of ALS are in agreement with prior reports of TDP-43-induced proteinopathy exhibiting reduced Futsch abundance at the NMJ due to diminished translation of its mRNA (Coyne et al., 2014). Therefore, multiple *Drosophila* models of ALS are associated with a decrease in microtubule stability stemming either from reduced Futsch abundance or due to hyperphosphorylation of the protein (Pennetta et al., 2002; Ratnaparkhi et al., 2008; Coyne et al., 2014). Interestingly, expression of a gain-of-function variant of *vapb* results in an increase in the number of synaptic boutons and number of Futsch loops (Sanhueza et al., 2014). Given the likelihood that VAPB^P58S^ acts through a dominant-negative mode of action (Ratnaparkhi et al., 2008), there appears to be a direct correlation between the functionality/dosage of VAPB, Futsch-dependent microtubule stability, and bouton number at the *Drosophila* NMJ (Pennetta et al., 2002; Sanhueza et al., 2014).

Ectopic expression of an overactive TRPV4 channel variant that elevates intracellular [Ca^2+^] leads to neuropathology via CaMKII hyperactivation (Woolums et al., 2020). Therefore, there is precedent for the notion that aberrant activation of CaMKII in motor neurons implies higher cytosolic [Ca^2+^]. Dampening sources of presynaptic Ca^2+^ elevation then would be expected to restore Ca^2+^ homeostasis, and thereby, ameliorate the consequences of CaMKII hyperactivation. Indeed, reduced expression of ER Ca^2+^ release channels in *Drosophila* neurons counteracts the phenotypes stemming from activating mutations in a voltage-gated Ca^2+^ channel (Brusich et al., 2018). We found that lowering the abundance of any one of the three presynaptic ER Ca^2+^ release channels — Iav, RyR, and IP_3_R — restored bouton morphology and number in neurons expressing *vapb^P58S^*. These data also point to the existence of a biphasic, bell-shaped relationship between presynaptic [Ca^2+^] and NMJ morphology, whereby either a supraphysiological increase or decrease in [Ca^2+^] elicits morphologically indistinguishable changes at the NMJ. While a decrease in presynaptic resting [Ca^2+^] results in Futsch hyperphosphorylation and microtubule destabilization due to diminished activity of the Ca^2+^/CaM-responsive phosphatase, calcineurin, increased cytosolic [Ca^2+^] also elicits Futsch hyperphosphorylation albeit via elevated CaMKII activity.

Given that the NMJ phenotypes in neurons expressing *vapb^P58S^* were suppressed by the loss of ER Ca^2+^ channels, which are needed for maintaining presynaptic resting [Ca^2+^] (Wong et al., 2014), we asked whether VAPB^P58S^ was triggering an increase in resting [Ca^2+^]. Higher presynaptic resting [Ca^2+^] augments the probability of SV release resulting in increased amplitudes of evoked post-synaptic potentials that is accompanied by paired-pulse depression (Wong et al., 2014). However, in agreement with prior reports (Chai et al., 2008), we found that expression of *vapb^P58S^* induced neither a change in the amplitude of evoked potentials nor the appearance of paired-pulse depression. Rather, *vapb^P58S^* NMJs exhibited paired-pulse facilitation, which in combination with the absence of changes in the amplitude of evoked potentials, was consistent with post-stimulus accumulation of residual Ca^2+^ due to delayed extrusion of presynaptic Ca^2+^. Indeed, direct examination of cytosolic [Ca^2+^] using GCaMP5G- tdTomato revealed significantly delayed Ca^2+^ extrusion in *vapb^P58S^*-expressing motor neurons. These data indicate that the aberrant activation of CaMKII in *vapb^P58S^*- expressing neurons arose from the accumulation of residual Ca^2+^ due to delayed extrusion.

### What could explain the defects in Ca^2+^ extrusion in neurons expressing vapb^P58S^?

The plasma membrane-resident Ca^2+^ ATPase, PMCA, has been shown to play a major role in the extrusion of presynaptic Ca^2+^ in *Drosophila* motor neurons (Ivannikov and Macleod, 2013; Rossano et al., 2013). Given the bioenergetic burden of powering PMCA, we postulated that a paucity of ATP could underlie the Ca^2+^ dyshomeostasis observed in *vapb^P58S^*-expressing neurons. In support of this model, mutated VAPB has been shown to disrupt interorganellar transfer of Ca^2+^ from the ER to mitochondria in both mammalian cells and *Drosophila* neurons (De Vos et al., 2012; Stoica et al., 2016; Gomez-Suaga et al., 2017; Smith et al., 2017; Xu et al., 2020; Wong et al., 2021). Given the role of matrix [Ca^2+^] in ATP production (Duchen, 1992; McCormack and Denton, 1993; Dumollard et al., 2004; Cárdenas et al., 2010; Ding et al., 2018), we asked whether expression of *vapb^P58S^* compromised mitochondrial ATP production. By examining the [ATP]/[ADP] ratio in live neurons, we found that ATP production was unable to match the demand of depolarization in *vapb^P58S^*-expressing neurons.

Whereas the [ATP]/[ADP] ratio in *vapb^WT^*-expressing neurons remained stable in response to depolarization, this ratio dropped precipitously in depolarized *vapb^P58S^* neurons. The situation in *vapb^WT^* neurons was akin to that in control *Drosophila* and mammalian neurons where the rates of ATP synthesis are exquisitely synchronized with its demand (Rangaraju et al., 2014; Wong et al., 2021). This synchrony was disrupted in *vapb^P58S^* neurons. The drop in the [ATP]/[ADP] ratio likely reflected a combination of diminished ATP synthesis and enhanced ATP consumption. Indeed, *in silico* modeling has predicted that mitochondrial dysfunction in ALS neurons is sufficient to induce a toxic self-reinforcing loop of increased ATP consumption and a progressive decrease in the availability of ATP (Le Masson et al., 2014). Taken together, our data are consistent with the model that the paucity of ATP levels in *vapb^P58S^*-expressing neurons during periods of neuronal activity result in diminished Ca^2+^ extrusion, which in-turn provokes hyperactivation of CaMKII and the attendant changes in NMJ morphology.

Maintenance of synaptic transmission at the *Drosophila* larval NMJs during periods of high-frequency stimulation depends upon a steady supply of ATP. A shortage of local ATP levels — for instance, due to an absence of presynaptic mitochondria in *drp1* mutants — results in the rundown of SV release in response to high-frequency stimulation (Verstreken et al., 2005). These rundowns occur due to the inability of the synapses to meet the energy demands of SV cycling and diminished recruitment of the reserve pool of SVs that are otherwise mobilized to replace the rapid depletion of the readily-releasable pool of vesicles (Verstreken et al., 2005). In either the *vapb^WT^*- or *vapb^P58S^*-expressing NMJs, high-frequency stimulation led to initial elevation of synaptic transmission. Superimposed over these liminal elevations were rundowns that became obvious after ∼5 minutes of stimulation, and were significantly greater in *vapb^P58S^*- expressing neurons. These data agree with enhanced rundown of SV release in motor neurons expressing *vapb^P58S^*. The magnitude of this phenotype, however, was smaller than that observed in the *drp1* mutants (Verstreken et al., 2005). This discrepancy may be explained by a steady-state decrease in ATP levels in *drp1* mutant neurons (Verstreken et al., 2005), while *vapb^P58S^*-expressing neurons exhibit ATP insufficiency only during periods of heightened activity (Fig. 5).

In addition to the early changes in synapse development and function, expression of *vapb^P58S^* also leads to adult-onset locomotor dysfunction and premature lethality (Wong et al., 2021). The findings presented here suggest that the neurodevelopmental and longevity phenotypes stem from partially overlapping, yet distinct, molecular pathways. While the NMJ phenotypes were suppressed by concomitant reductions in the abundance of Iav and RyR, or by the expression of the CaMKII inhibitory peptide, none of these manipulations influenced the effects of VAPB^P58S^ on adult lifespan. Only with the knockdown of the gene encoding IP_3_Rs do we observe suppression of both the NMJ development phenotypes and adult lethality, though the former involves the attenuation of CaMKII while the latter, as we showed previously, stems from the prevention of endolysosomal [Ca^2+^] overload (Wong et al., 2021).

## Acknowledgements

We thank the Bloomington *Drosophil*a Stock Center for fly stocks. We also thank Yufang Chao for technical help. Confocal and live cell microscopy was performed at the Center for Advanced Microscopy, Department of Integrative Biology & Pharmacology at McGovern Medical School, UTHealth. This work was supported by the NIH grants RF1AG069076, RF1AG067414, and R21AG072176 (all to K.V.).

## Author Contributions

C.W., N.E.K., K.L.T, and K.V. conducted the experiments. C.W., N.E.K., K.L.T, and K.V. analyzed the data. C.W., N.E.K., H.J.B., and K.V. conceived the experiments, and wrote the manuscript.

## Materials and Methods

### Drosophila husbandry

Flies were reared at 21 °C on standard fly food (1 L of food contained: 95 g agar, 275 g Brewer’s yeast, 520 g of cornmeal, 110 g of sugar, 45 g of propionic acid, and 36 g of Tegosept) unless otherwise stated. The following fly lines were obtained from Bloomington *Drosophila* Stock Center: *ok371-GAL4* and *d42-GAL4* (Parkes et al., 1998; Mahr and Aberle, 2006), *RyR^16^* (Sullivan et al., 2000), *UAS-Dcr1* (Dietzl et al., 2007), *UAS-CKII-I.Ala* (*UAS-CKII^ala^*) (Joiner MlA and Griffith, 1997), and *UAS-itpr^RNAi^* (*TRiP.JF01957*) (Wong et al., 2021). Other strains used in the study were: *UAS-vapb^WT^* and *UAS-vapb^P58S^* (Tsuda et al., 2008), *UAS-GCaMP5G-tdTomato* (Wong et al., 2021), *Canton-S*, *UAS-PercevalHR*(Wong et al., 2021), *iav^1^* (also called *iav^hypoB-1^*) (Wong et al., 2014), and *UAS-iav* (Wong et al., 2014)*. Canton-S* flies were used as the wild-type controls in the various crosses.

### NMJ immunohistochemistry and confocal microscopy

Dissection and immunostaining of NMJ was performed as described previously (Wong et al., 2014, 2015). Briefly, wandering 3^rd^ instar larvae were filleted in ice-cold PBS to remove all visceral organs except the brain and nerves. The fillets were fixed in 4% paraformaldehyde in PBS for 30 minutes. The fixed fillets were washed with 0.1% Triton X-100 in PBS before incubation with primary antibodies overnight at 4°C. Primary antibody dilutions were 1:100 mouse anti-discs large (DLG) and 1:50 mouse anti-Futsch. The monoclonal antibodies against DLG and Futsch were obtained from the Developmental Studies Hybridoma Bank developed under the auspices of the NICHD and maintained by The University of Iowa, Department of Biology, Iowa City, IA 52242. The samples were then washed and probed with 1:200 FITC-conjugated anti-HRP (Jackson ImmunoResearch) and Alexa Fluor 568-conjugated anti-mouse secondary antibodies (ThermoFisher) at room temperature for 1.5 hours, and then mounted on glass slides with Vectashield (Vector Labs). Confocal images were obtained using a Nikon A1 Confocal Laser Microscope System (Nikon). For NMJ bouton counting, a 60x oil objective was used to focus on the NMJs on abdominal segment 3.

### NMJ electrophysiology

Wandering 3^rd^ instar larvae were dissected in ice-cold HL-3 (70 mM NaCl, 5 mM KCl, 20 mM MgCl_2_.6H_2_O, 10 mM NaHCO_3_, 115 mM sucrose, 5 mM trehalose, and 5 mM HEPES; pH 7.2) and rinsed with HL-3 containing 0.5mM Ca^2+^. Recordings were made from body-wall muscle 6 (abdominal segment 3) with sharp electrodes filled with a 2:1 mixture of 2 M potassium acetate and 2 M potassium chloride. Data were collected from samples with resting membrane potential below −60 mV. EJPs were evoked by directly stimulating the A3 hemisegmental nerve through a glass capillary electrode (internal diameter, ∼10μm) at 0.2 Hz. Stimulus pulses were generated by pClamp 10 software (Axon Instruments Inc). The applied currents were 6 μA ± 3 with fixed stimulus duration at 0.3 ms. Twenty to thirty evoked EJPs were recorded for each muscle for analysis. Miniature EJP (mEJP) events were collected for 5 min. Both EJPs and mEJPs were amplified with an Axoclamp 900A amplifier (Axon Instruments, Foster City, CA) under bridge mode, filtered at 10 kHz and digitized at 10 kHz (EJPs) and 40 kHz (mEJPs) with pClamp 10. Experiments were performed at 21 °C. EJPs and paired-pulse stimulation were analyzed with pClamp 8.0 software (Axon Instruments). The mEJPs were analyzed using the Mini Analysis Program (Synaptosoft Inc., Decatur, GA). The EJPs paired-pulse amplitudes were corrected by nonlinear summation (Feeney et al., 1998). Paired-pulse ratio was calculated as the ratio of 2^nd^ to 1^st^ peak. The quantal content of evoked release was calculated from individual muscles by the ratio of the average EJP amplitude over the average mEJP amplitude. For high-frequency stimulation, nerves were stimulated for 10 minutes at 10 Hz. By fitting the decay portions of each high-frequency trace to a 1^st^-order exponential function, we obtained the rate-constant of decays. Rundowns were extracted from the rate-constants of decay. CI of the calculated rundowns were determined by fitting the data calculated rundowns to 1^st^- order exponential functions in Prism 8. *P*-values for the confidence intervals (CI) were calculated as described (Altman and Bland, 2011).

### Dissociation of *Drosophila* neurons

We dissociated primary motor neurons from *Drosophila* as described previously (Wong et al., 2021). Briefly, the exterior of wandering 3^rd^ instar larvae was sterilized by brief submersion in ethanol, and then washed with sterilized H_2_O before dissection in filtered Schneider’s medium (S0146; Sigma-Aldrich) containing 10% FBS, antibiotic/antimycotic solution (A5955; Sigma-Aldrich) and 50 μg/ml of insulin (I6634; Sigma-Aldrich). Brains dissected from these larvae were washed in separate wells containing filtered Schneider’s medium before being transferred to a filtered HL-3 solution (70 mM NaCl, 5 mM KCl, 1 mM CaCl_2_, 20 mM MgCl_2_, 10 mM NaHCO_3_, 115 mM sucrose, 5 mM trehalose, and 5 mM HEPES) supplemented with 0.423 mM L-cysteine (Calbiochem) and 5 U/mL papain (Worthington) (Note: After L-cysteine addition but before papain addition, the pH of the solution was recalibrated to 7.4). The brains were then enzymatically digested in the papain solution for 20 minutes before transfer to a 1.5 mL tube containing 1 mL of filtered Schneider’s medium. Cells were centrifuged at 100G for 1 minute prior to decantation of Schneider’s medium. The solution was replaced with 1 mL of fresh filtered Schneider’s medium. This process was repeated twice before neurons were dissociated by pipetting repeatedly until the solution was homogeneous. The solution with dissociated neurons was then placed on 35 mm glass bottom dishes (D35-10-0-N; Cellvis) that had been coated with concanavalin A (C2010; Sigma-Aldrich). Cells were cultured in Schneider’s medium supplemented with 10% FBS, antibiotic/antimycotic solution (A5955; Sigma-Aldrich) and 50 μg/mL of insulin (I6634; Sigma-Aldrich) at room temperature in a humidified container at room temperature. After each day in culture, cells were washed twice with PBS to remove any yeast contamination or debris remaining from dissociation.

### Live-cell imaging of fly primary neurons

Live imaging of dissociated neurons was performed as described previously (Wong et al., 2021). Briefly, culture media on plates of dissociated neurons was first replaced with HL-3 (70 mM NaCl, 5 mM KCl, 1 mM CaCl_2_, 20 mM MgCl_2_, 10 mM NaHCO_3_, 115 mM sucrose, 5 mM trehalose, and 5 mM HEPES; pH 7.2, room temperature). For measurements of cytosolic Ca^2+^, GCaMP5G and tdTomato were sequentially excited at 488 nm and 561 nm respectively by an A1 laser confocal microscope with a 40x objective (Nikon). Emission signals at 525 nm and 595 nm were recorded. Backgrounds were measured from a cell-free ROI. Baselines were established for 1 minute before addition of muscarine (1 mM). Cytosolic Ca^2+^ transients were evoked by store-depletion with the SERCA inhibitor, thapsigargin, as described. Amplitudes of the GCaMP5G/tdTomato ratio represented cytosolic free [Ca^2+^]. >50 cells from a minimum of 5 independently conducted experiments for each condition were used for the calculations.

PercevalHR signals were recorded by measuring the ratio of fluorescence emissions at 525 nm sequentially excited at 487.5 nm and 407.8 nm. An A1R laser confocal microscope with 40x objective (Nikon) was used for measurement. Background emission signals were measured from a cell-free ROI. Baselines were established for 1 minute before addition of muscarine (1 mM). For cells subjected to depolarization, the bath was replaced with high [K^+^] (52 mM) HL-3. Muscarine (1 mM), 2-DG (10 mM), and oligomycin A (10μM) were added as needed and signals were recorded. Amplitudes of the emission ratio represented the cytosolic [ATP]/[ADP] ratio (Tantama et al., 2013; Wong et al., 2021). Data were quantified as change in the PercevalHR ratio over the baseline prior to the addition of the drug. For this quantification, we used custom R code to calculate the integrated change (i.e., area under the curve using the AUC function in R) per unit time. >80 cells from a minimum of 3-4 independently conducted experiments for each condition were used for the calculations.

### Analysis of fly lifespan

Newly eclosed adult flies were collected and transferred to vials containing standard fly food (≤ 15 flies per vial). Flies were kept at room temperature (∼21°C) and transferred to new vials twice per week. Dead flies at the bottom of the old vials were counted after each transfer until all the animals in a cohort died.

### Statistical analyses

We used either a parametric or a nonparametric test of statistical significance on the basis of whether the data were normally distributed. Multiple comparisons were made by ANOVA. R, Excel (Microsoft), and Prism 8 (GraphPad) were used for statistical analyses. We used also used Prism for curve fitting. Custom R code used for quantifying data is available upon request. Statistical significance was defined as a *P <* 0.05. *P*-values were shown on the figures as *asterisks*: *, *P* < 0.05; **, *P* < 0.01; ***, *P* < 0.001; ****, *P* < 0.0001. Lifespan (Kaplan-Meier) curves were generated using Prism 8. We used the log-rank (Mantel-Cox) test to determine *P*-values.

